# Plants with self-sustained luminescence

**DOI:** 10.1101/809376

**Authors:** Tatiana Mitiouchkina, Alexander S. Mishin, Louisa Gonzalez Somermeyer, Nadezhda M. Markina, Tatiana V. Chepurnyh, Elena B. Guglya, Tatiana A. Karataeva, Kseniia A. Palkina, Ekaterina S. Shakhova, Liliia I. Fakhranurova, Sofia V. Chekova, Aleksandra S. Tsarkova, Yaroslav V. Golubev, Vadim V. Negrebetsky, Sergey A. Dolgushin, Pavel V. Shalaev, Olesya A. Melnik, Victoria O. Shipunova, Sergey M. Deyev, Andrey I. Bubyrev, Alexander S. Pushin, Vladimir V. Choob, Sergey V. Dolgov, Fyodor A. Kondrashov, Ilia V. Yampolsky, Karen S. Sarkisyan

**Author notes:** equal contribution (Tatiana Mitiouchkina, Alexander S. Mishin, Louisa Gonzalez Somermeyer, Nadezhda M. Markina, Ilia V. Yampolsky, Karen S. Sarkisyan).

## Abstract

In contrast to fluorescent proteins, light emission from luciferase reporters requires exogenous addition of a luciferin substrate. Bacterial bioluminescence has been the single exception, where an operon of five genes is sufficient to produce light autonomously. Although commonly used in prokaryotic hosts, toxicity of the aldehyde substrate has limited its use in eukaryotes^1^. Here we demonstrate autonomous luminescence in a multicellular eukaryotic organism by incorporating a recently discovered fungal bioluminescent system^2^ into tobacco plants. We monitored these light-emitting plants from germination to flowering, observing temporal and spatial patterns of luminescence across time scales from seconds to months. The dynamic patterns of luminescence reflected progression through developmental stages, circadian oscillations, transport, and response to injuries. As with other fluorescent and luminescent reporters, we anticipate that this system will be further engineered for varied purposes, especially where exogenous addition of substrate is undesirable.

## Introduction

Using optical reporters for imaging in model organisms is often limited by light scattering and absorption. Fluorescent reporters in particular are further hampered by endogenous fluorophores, especially in plants which harbor an abundance of fluorescent compounds. By not requiring light excitation, bioluminescence circumvents autofluorescence and abates light scattering and absorption. For these reasons, bioluminescence is commonly preferred in animal models where it provides greater detection sensitivity and quantitation. These benefits would be augmented in plants by their more challenging optical characteristics, except that the requirement for an exogenously supplied substrate has been a substantial obstacle. A genetically encodable autoluminescence pathway in plants would thus have considerable potential, with the fungal bioluminescent system being a suitable candidate for the development of such technology.

Bioluminescent fungi emit light in the green part of the visible spectrum, fitting well into the optical transparency window of pigmented plant tissues (**Figure 1**, **inset**). In addition, the biochemical pathway leading to light production in fungi is well suited for integration into plant metabolism. Termed the caffeic acid cycle, it is a short metabolic route stemming from caffeic acid, a ubiquitous compound in vascular plants (**Figure 1**). Caffeic acid is among the core metabolites of the phenylpropanoid pathway leading to lignols, flavonoids, anthocyanins, stilbenes, coumarins and numerous other classes of phenolic compounds ^3^(**Figure 1**).

**Figure 1.**
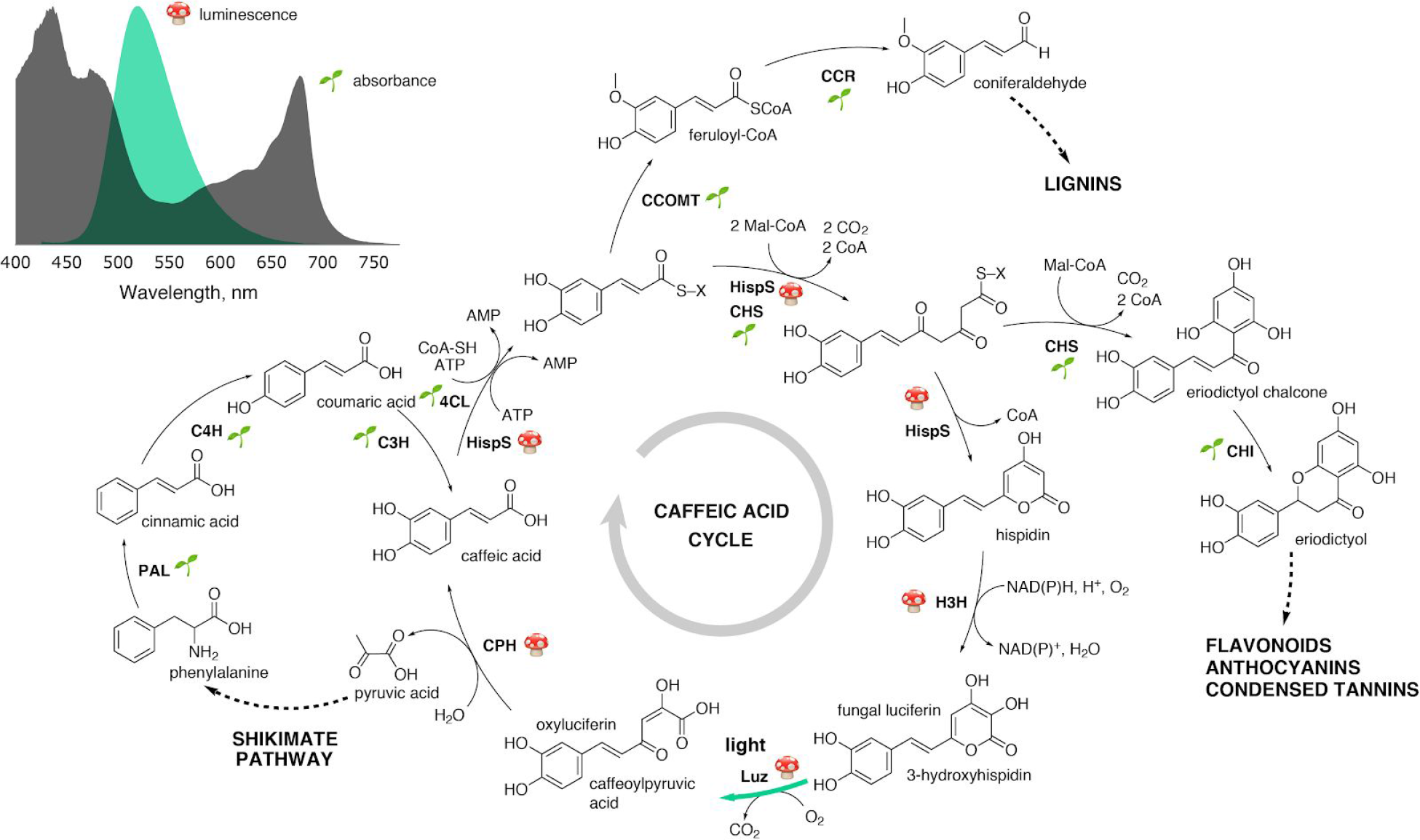
Spectral and biochemical characteristics of the fungal bioluminescence system relevant for engineering glowing plants. The caffeic acid cycle shares metabolites with some of the major plant biosynthetic pathways. The fungal or plant origin of enzymes is indicated with the symbols 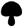and 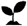, respectively. Abbreviations: 4CL — 4-coumarate:CoA ligase; C3H — p-coumaric acid 3-hydroxylase; C4H — cinnamic acid 4-hydroxylase; CCOMT — caffeoyl-CoA 3-O-methyltransferase; CCR — cinnamoyl-CoA reductase; CHI — chalcone isomerase; CHS — chalcone synthase; CPH — putative caffeoylpyruvate hydrolase; H3H — hispidin-3-hydroxylase; HispS — hispidin synthase; Luz — luciferase; PAL — phenylalanine ammonia-lyase. **Inset**: spectrum of fungal bioluminescence (*Neonothopanus nambi*, in green) overlaid onto the absorbance spectrum of plant leaves (*Nicotiana tabacum*, in dark gray).

## Results

We engineered autonomously glowing tobacco plants by constitutively expressing enzymes of the caffeic acid cycle ^2^ derived from the fungus *Neonothopanus nambi* (**Supplementary Note 1**). The pathway is driven by the activities of four enzymes: luciferase Luz; two enzymes of luciferin biosynthesis, hispidin synthase HispS and hispidin-3-hydroxylase H3H; and a putative oxyluciferin recycling enzyme CPH (**Figure 1**). Transgenic plants expressing these genes emitted light at all developmental stages, without the need for externally supplied substrates and with sufficient brightness to be visible to the naked eye. We were able to capture detailed images on consumer-grade cameras and even on smartphones, with exposure times as low as 5–30 seconds (**Supplementary Fig. 1–3**, **Figure 2**).

**Figure 2.**
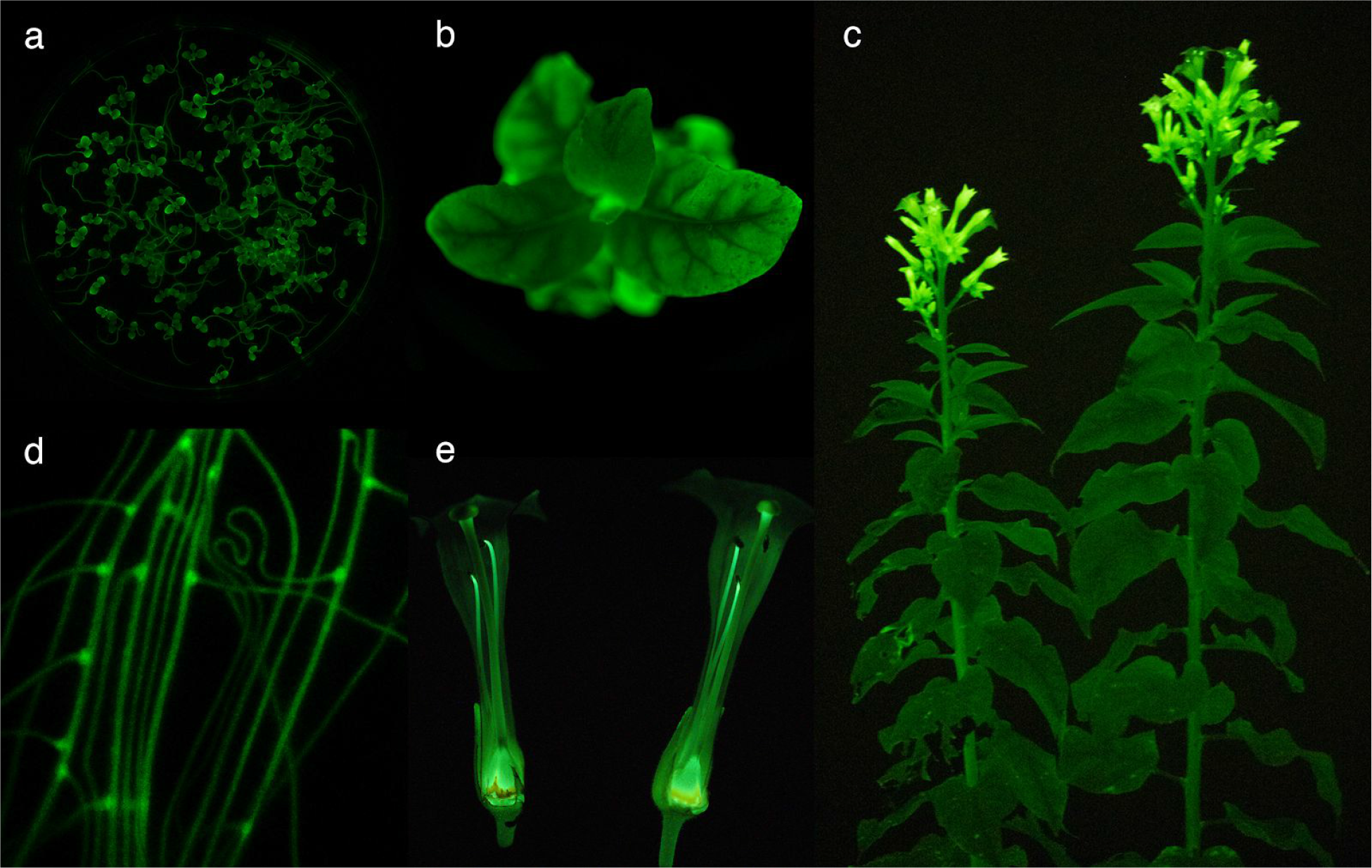
Bioluminescent plants at various stages of development. Light emission from *N. tabacum* plants at germination (**a**), vegetative (**b**) and flowering (**c**) stages. Roots (**d**) and cross section of flowers (**e**). Photos were captured on Sony Alpha ILCE-7M3 (see **Methods**).

The luminescence of engineered plants showed dependence on the developmental stage and on circadian rhythms, and changed in response to physiological stresses. Seeds of transgenic tobacco lines started glowing upon germination (**Figure 2a**, **Supplementary Video 1**), with major light sources being the tips of cotyledons and roots. The roots glowed particularly bright at branching points (**Figure 2d**), often hours before lateral root initiation became evident in ambient light (**Supplementary Videos 2–3**). As plants developed, luminescence remained unevenly distributed within the plant, being brightest at the transition zone between the root and the stem. The overall phenotype, leaf color, flowering time, and seed germination were not noticeably different from those of the wild-type tobacco, suggesting that unlike bacterial bioluminescence, high expression of the caffeic acid cycle is not toxic and does not impose a significant burden on plant growth (**Supplementary Figures 1-2; Supplementary Note 2**).

In young plants, the brightest parts of the shoot were the terminal and axillary buds and the upper part of the stem. As plants matured, the older parts of the shoot dimmed (**Supplementary Video 4**). At the flowering stage, the glow of flower buds surpassed the luminescence from other parts of the plant. Luminescence was strikingly brighter in the petals and particularly the ovary, and apical portions of the style and stamen filaments (**Figure 2c, 2e**, **Supplementary Figure 4**). Notably, the distribution of luminescence resembled reported expression patterns for phenylalanine ammonia-lyase ^4^, an entry point of the phenylpropanoid pathway (**Supplementary Note 3**). These observations suggest that light intensity is linked to metabolic activity and particularly may reflect the availability of caffeic acid.

To identify metabolites that may be limiting light emission, we infused leaves of glowing plants with all-but-one mixtures of hispidin precursors, as well as with each precursor individually. As *Nicotiana tabacum* leaves did not retain the precursors at the place of injection, we created a similar glowing strain of *Nicotiana benthamiana* for these experiments. Infusion of leaves with malonyl-CoA, CoA or ATP individually or as a mixture did not increase light emission. By contrast, solutions containing caffeic acid induced an increase in luminescence (**Supplementary Figure 5**). This suggests that caffeic acid limits hispidin biosynthesis. Limitation by the other precursors cannot be excluded (**Supplementary Note 4**) but since they all participate in the phenylpropanoid pathway, light emission should be effectively coupled to the activity of this pathway.

If light emission is linked to phenylpropanoid metabolism, luminescence is expected to become brighter under conditions known to activate production of phenylpropanoids, such as leaf injury ^5,6^ and removal of apical shoots ^7^. The concentration of phenylpropanoids is also known to increase during flower development ^8^, senescence-related nutrient remobilization in leaves ^9,10^ or after treatment with methyl jasmonate ^11,12^ or ethylene ^13^. We evaluated light emission under these conditions to further explore linkage of bioluminescence to phenylpropanoid metabolism.

Indeed, in injured leaves, we observed a sustained increase in light emission at the injury site. We have also registered luminescence spreading from the injury site by small veins with the velocity of about 2 μm/sec (**Supplementary Figure 6, Supplementary Video 6**). Apical shoot removal also resulted in sustained bright luminescence in lateral shoots proximal to the cut site (**Supplementary Figure 8, Supplementary Video 7**). Aging leaves, reported to have gradually reducing caffeic acid content until late senescence ^14^, generally exhibited decreased light emission. Nevertheless, some leaves displayed waves of intense light emission during the final stages of senescence (**Supplementary Video 4**), possibly reflecting age-related nutrient remobilization. Finally, plants treated with methyl jasmonate or ethylene responded with dramatically increased luminescence throughout the plant (**Figure 3ab**). Thus, in all cases, time-lapse imaging revealed the expected increase in light emission.

**Figure 3.**
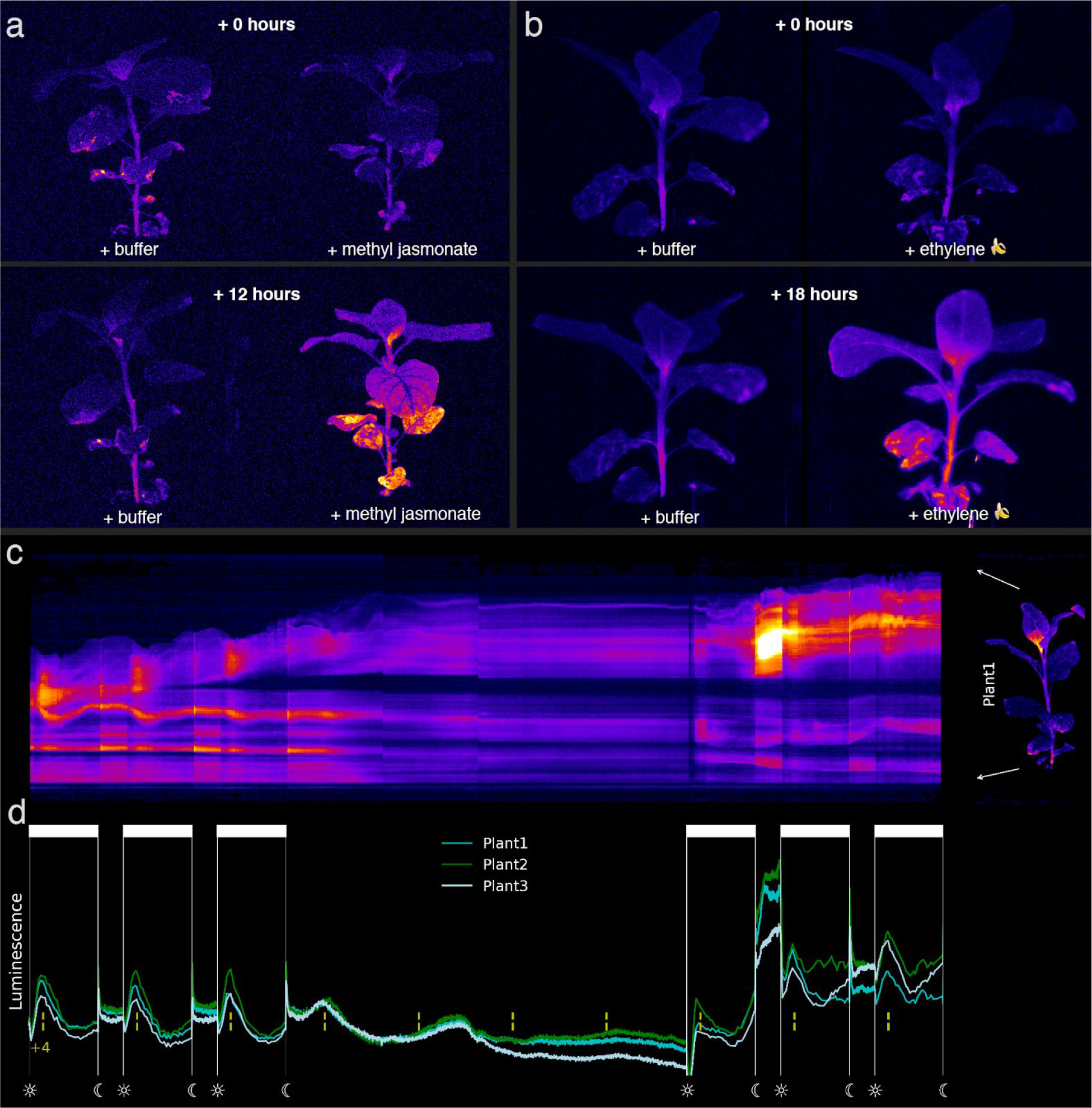
Dynamics of luminescence in living plants. **A**. Photo of transgenic *Nicotiana tabacum* plant sprayed with 5 mM methyl jasmonate solution or buffer (control) before and after treatment. **B.** Photo of transgenic *Nicotiana tabacum* plant incubated in a closed vial with banana skin. **C**. Circadian oscillations of luminescence. Photos of three plants were being captured constantly for ten days, in normal light conditions (17.5 h day; days 1-3 and 8-10) or in constant darkness (days 4-7). Each image of Plant 1 was compressed into a single vertical line of pixels to create the kymogram **C**. The mean brightness of plants 1-3 is displayed on the graph **D**.

Luminescence in living plants was quite dynamic (**Supplementary Video 4**). Throughout their lifespan, plants displayed circadian oscillations of light emission, from germinating seeds to flowering plants ^8,15^ (**Figure 3c**, **Supplementary Videos 1, 4, 8**). Often, bright bioluminescence appeared in leaves as flickering dots or ovals with a diameter of a few millimeters. This flickering pattern was especially evident in young leaves, particularly those near brightly glowing pruning-induced axillary shoots. Imaging the lower side (abaxial surface) of these leaves revealed luminescence dynamics suggesting intercellular transfer events possibly associated with phloem transport (**Supplementary Video 7**). Finally, in young leaves, we observed dynamic waves of bright luminescence spreading throughout the blades during the day (**Supplementary Figure 8, Supplementary Video 9**). Homogeneous distribution of signal, not associated with the leaf vascular system, suggests a transport-independent mechanism behind the increase in light emission and may correspond to previously reported spatiotemporal waves of gene expression ^16^.

## Discussion

The caffeic acid cycle from fungi is one example of the many distinct luminescent systems found among evolutionary distant organisms, ranging from bacteria to protists to fungi and animals ^17^. Biochemical characterization of these systems has demonstrated that light emission is achievable via multiple metabolic routes ^18^. Several luciferases have been used as reporters of biomolecular processes, most notably firefly luciferase, various marine luciferases, such as Renilla, Gaussia, Oplophorus and others, and bacterial luciferase ^19^. They have proven to be enormously useful primarily because luminescence can be readily quantified with high sensitivity and precision.

However, the requirement for exogenously added luciferin can be cumbersome, especially for complex multicellular organisms where substrate permeability and tissue distribution can be challenging. While it has long been desirable to achieve endogenous production of luciferin, this has remained elusive with the exception of the operon associated with bacterial bioluminescence. Furthermore, by examination of luciferin structures, it is not apparent whether their biosynthesis would be readily compatible with relevant plant or animal models. For example, firefly luciferin is likely produced through condensation with D-cysteine, an inverse enantiomer of the native amino acids in proteins ^20^. Thus, bacterial bioluminescence has previously been the only prospect for achieving autonomous luminescence. Yet, more than three decades since the operon was discovered, this system has not become widely used in eukaryotic hosts.

Discovery of the enzymes driving fungal bioluminescence provides a new opportunity for producing autonomous luminescence in higher organisms. This opportunity may be greatest for vascular plants due to the fortuitous commonality of metabolites. Luminescence is not known to occur naturally in any plants ^17^. Fungi, by comprising a separate living kingdom, are only distantly related. Yet, the fungal metabolites required for hispidin biosynthesis (caffeic acid, malonyl-CoA, and ATP) are also components of the phenylpropanoid pathway in plants. Moreover, low levels of hispidin and related styrylpyrones are present in some vascular plants (**Supplementary Table 1; Supplementary Note 1**). Only the substrate and product directly utilized by the fungal luciferase are apparently unique. Although caffeic acid is not native to animals, autonomous luminescence is possible by including two additional enzymes needed for its biosynthesis from tyrosine — tyrosine ammonia lyase TAL and coumarate 3-hydroxylase C3H, or their functional equivalents (**Supplementary Figure 12**) ^21^.

The diversity of fluorescent compounds in plants interferes with detection of GFP and related fluorescent proteins ^22,23^. Besides the strong fluorescence of chlorophyll extending beyond 640 nm, compounds emiting in the 500–600 nm range include phenols, flavins, polyacetylene, and isoquinoline. As plants lack intrinsic luminescence, luciferase reporters can be detected at much lower expression levels. Nonetheless, in this initial feasibility study, we estimate the glowing plants to be more than an order of magnitude brighter than previously reported using bacterial bioluminescence ^24^. We anticipate that light emission could be further increased through optimizing catalytic efficacy of the fungal enzymes in plants ^25,26^, improving metabolic channelling between the enzymes ^27^, and enhancing precursors availability through metabolic engineering ^28^.

Rapid dynamic measurements are possible with bioluminescence since light emission arises from an enzymatic process. Thus, light intensity responds quickly to perturbations of the catalytic steady state. This is precluded with fluorescent proteins due to the long persistence of their chomophore. Benefiting from this, we have observed waves of phenylpropanoid activity spreading from injury sites, possibly dynamics of phloem unloading in sink leaves, and senescence-related metabolic events in leaves. We believe our direct observations of these dynamic processes are the first reported. As configured in this initial study, light emission was coupled to phenylpropanoid metabolism. However, through various strategies involving energy transfer to fluorophores, ratiometric measurements could allow isolation of the reporter response to other biomolecular processes, including quantitative measurements of gene activity, cellular signalling and other physiological activities ^29,30^. In particular, the results obtained by Khakhar et al. in parallel to this work showed robust performance of the fungal bioluminescent system as a reporter of gene expression in plants ^31^.

To recapitulate fungal bioluminescence in the tobacco plants, we had transfected all four of the genes represented in the caffeic acid cycle. However, expression of only three enzymes (nnHispS, nnH3H, and nnLuz) was sufficient to achieve autonomous luminescence. Additional expression of the putative luciferin recycling enzyme (nnCPH) did not increase brightness in the plants (**Supplementary Note 1**). Forming gene fusions may also reduce the number of necessary reading frames, thereby further simplifying the genetic requirements. Greater understanding of the structures and biochemical properties of the fungal enzymes will likely produce additional simplifications. Thus, with some optimization, we anticipate that integrating fungal bioluminescence into plants should be relatively straightforward. By enabling autonomous light emission with this metabolically compatible means, dynamic processes can be monitored continuously in a variety of living plants. This may include dynamics associated with plant development and pathogenesis, responses to environmental conditions, and interventions through synthetic chemicals. Rapid screening methods should also be supported by the efficiency of acquiring luminescent data. Finally, glowing plants are aesthetically pleasant, offering new possibilities for artistic expression using luminescence.

## Methods

### Plasmid assembly

Coding sequences of the nnLuz, nnHispS, nnH3H and nnCPH genes were optimized for expression in *Nicotiana tabacum* and ordered synthetically from Evrogen (Moscow, Russia). Genes were later assembled into vectors for Golden Gate cloning following MoClo standard described in ref. ^32^. Each transcriptional unit contained 35s promoter, a fungal gene and *ocs* terminator. Final vectors for plant transformation contained kanamycin resistance cassette as well as transcriptional units for expression of fungal genes.

All Golden Gate cloning reactions were performed in 1X T4 ligase buffer (ThermoFisher) containing 10U of T4 ligase, 20U of either BsaI or BpiI (ThermoFisher), and 100ng of DNA of each DNA part. Golden Gate reactions were performed according to “troubleshooting” cycling conditions described in ref. ^33^: 25 cycles between 37°C and 16°C (90 sec at 37°C, 180 sec at 16°C), then 5 min at 50°C and 10 min at 80°C.

### Agrobacterium-mediated transformation

Assembled plasmids were transferred into *Agrobacterium tumefaciens* strain AGL0 ^34^. Bacteria were grown on a shaker overnight at 28°C in LB medium supplemented with 25 mg/l rifampicin and 50 mg/l kanamycin. Bacterial cultures were diluted in liquid Murashige and Skoog (MS) medium to the optical density of 0.6 at 600 nm.

Leaf explants used for transformation experiments were cut from two-week-old tobacco plants (*Nicotiana tabacum* cv. Petit Havana SR1, *Nicotiana benthamiana*) and incubated with bacterial culture for 20 minutes. Leaf explants were then placed onto filter paper overlaid on MS medium (MS salts, MS vitamin, 30 g/L sucrose, 8 g/L agar, pH 5.8) supplemented with 1 mg/L 6-benzylaminopurine and 0.1 mg/L indolyl acetic acid. Two days after inoculation, explants were transferred to the same medium supplemented with 500 mg/L cefotaxime and 75 mg/L kanamycin. Regeneration shoot were cut and grown on MS medium with antibiotics.

### Molecular analysis of transgenic plants

Genomic DNA was extracted from young leaves of greenhouse-grown plantlets using cetyltrimethylammonium bromide method ^35^. The presence each of the transferred genes was confirmed by PCR with gene-specific primers (**Supplementary Table 2**).

For Southern blot, 30 μg of plant genomic DNA was digested overnight at 37°C by 100U of EcoRV, a restriction enzyme that cuts T-DNA constructs used in this study at a single position inside nnHispS coding region. After gel electrophoresis, digestion products were transferred onto Amersham Hybond-N+ membrane (GE Healthcare, UK) and immobilized. The DNA probe was constructed by PCR using cloned synthetic nnLuz gene as the template and nnLuz-specific primers listed in **Table 1**. Probe DNA was labeled with alkaline phosphatase using the AlkPhos Direct Labeling Kit (GE Healthcare, UK). Prehybridization, hybridization (overnight at 60°C) with alkaline phosphatase-labeled probe, and subsequent washings of the membrane were carried out according to the AlkPhos Direct Labeling Kit protocol. Detection was performed using Amersham CDP-Star detection reagent following the manufacturer’s protocol (GE Healthcare, UK). The signal from the membrane was accumulated on X-ray film (XBE blue sensitive, Retina, USA) in film cassette at room temperature for 24 hours. X-ray films were scanned on Amersham imager 600 (GE Healthcare Life Sciences, Japan).

### Plant growth conditions

Tobacco plants were propagated on Murashige and Skoog (MS) medium supplemented with 30 g/l sucrose and 0.8 w/v agar (Panreac, Spain). *In vitro* cultures were incubated at 24±1°C with 12-16-day photoperiod, with mixed cool white and red light (Cool White and Grolux fluorescent lamps) at light intensity 40 μmol / sec*m^2^. After root development, plantlets were transferred to 9 cm pots with sterilized soil (1:3 w/w mixture of sand and peat). Potted plants were placed in the greenhouse at 22±2°C under neutral day conditions (12h light / 12h dark; 150 μmol m-2s-1) and 75% relative humidity.

### Plant imaging setup with photo cameras

We used Sony Alpha ILCE-7M3 camera to capture all photos and videos presented in this paper, except those taken on a smartphone (**Supplementary Fig. 3**) and a long-term time-lapse filmed on Nikon D800 (**Supplementary Video 4**). Depending on the experimental setup, lens aperture and other considerations, a range of ISO values from 6400 to 40000 was used, with exposure times from 5 seconds (leaf injury) to 20 minutes (root microscopy). Most of the photos were captured with 30-second exposure time.

We used SEL50M28 lens (Sony, f/2.8), or 35mm T1.5 ED AS UMC VDSLR lens (Samyang, ~f/1.4). Long-term timelapse of growing tobacco plants (**Figure 3c**, **Supplementary Video 4**) was captured with Nikon D800 camera and Sigma AF 35mm f/1.4 DG HSM Art at ISO 8063 and 30-second shutter speed. Root microscopy was performed with Sony Alpha ILCE-7M3 camera with Meiji MA833 U. Plan 20X Objective lens was mounted on the camera via a custom-made adaptor.

The photos were then processed in the following way. First, a raw photo obtained in the dark with the same settings was per-channel subtracted from a raw photo of plants (LibRaw version 0.19.2, 4channels tool) to remove hot pixels and reduce noise. Then, an ImageJ plugin was applied (https://imagej.nih.gov/ij/source/ij/plugin/filter/RankFilters.java) to remove outliers (hot pixels). For most photos, only the green channels (G and G2) were kept in the final image.

### Plant imaging on a smartphone

We used smartphone Huawei P30 Pro for photography. To capture the photo displayed on **Supplementary Figure 3**, we used the following settings: 30 sec exposure time, ISO 6400, aperture 1.6.

### Absorption spectra of tobacco leaves

The leaves from adult wild type Nicotiana tabacum plants were collected and measured directly by spectrophotometer Cary 100 Bio (Varian).

### Imaging of leaf injuries

Plants were cultivated in greenhouse for six weeks. Leaves of *Nicotiana tabacum* were wounded with a blade causing cut across the midvein.

### Treatment with methyl jasmonate

3 week old transgenic bioluminescent *Nicotiana tabacum* plants were treated with methyl jasmonate (5 mM in 10 mM MES buffer pH 7.0) by spraying. Control plants were treated with buffer (10 mM MES buffer pH 7.0). Plants were then imaged in closed glass jars for 3 days in the dark.

### Incubation with banana skin

3 week old transgenic bioluminescent *Nicotiana tabacum* plants were imaged with ripe banana skin in closed glass jars for 24 hours.

### Quantitative PCR

In experiments aimed to determine whether expression of *luz* gene oscillates during the day, we collected the third leaves from twenty seven 25-day-old transgenic glowing plants. The leaves were collected with three-hour intervals during 24 hours, and to reduce the biological noise, leaves from three plants were combined. All leaves were flash-freezed in liquid nitrogen and homogenized for RNA extraction with TRIzol kit (Thermo Fisher Scientific, USA). Synthesis of the first cDNA strand was carried out with MMLV kit (Evrogen, Russia). Quantitative PCR was performed with qPCRmix-HS SYBR+LowROX kit (Evrogen, Russia) on 7500 Real-Time PCR machine (Applied Biosystems, USA) with primers annealing at *nnluz* transcript: GGACCAGGAGTCCCAGGC and CTTGGCATTTTCGACAATCTTA.

### Infiltration of *Nicotiana benthamiana* leaves with hispidin precursors

For experiments with infiltration of transgenic *Nicotiana benthamiana* leaves, we prepared 100 uM solutions of caffeic acid, malonyl-CoA, ATP and coenzyme A in 10 mM MES buffer (pH 7.0). We also prepared 100 uM mixtures of these compounds in the same buffer: *Mix 1*, *full* (caffeic acid, malonyl-CoA, CoA, ATP), *Mix 2 without caffeic acid* (malonyl-CoA, CoA, ATP), *Mix 3 without malonyl-CoA* (caffeic acid, CoA, ATP), and *Mix 4 without CoA* (caffeic acid, malonyl-CoA, ATP), *Mix 5 without ATP* (caffeic acid, malonyl-CoA, CoA). The solutions were injected into the blades of cut *Nicotiana benthamiana* leaves, and leaves were imaged for 15 min following injections. The analysis of the frame at one minute after the injection is presented on Supplementary Figure 5.

### LC-MS/MS analysis

Analytical standard (≥98.0) caffeic acid and acetic acid were purchased from Sigma-Aldrich. Hispidin was synthesised by Planta (≥95.0 %). HPLC-grade-acetonitrile was purchased from J.T. Baker. Deionized water was obtained from a Milli-Q System (USA).

We analysed several groups of samples: leaves and flowers of the wild type *Nicotiana tabacum* (NT000) and two transgenic lines of plants (NT001, NT078). Immediately after collection, the samples were frozen in liquid nitrogen and manually ground in a mortar. To reduce biological variability, we mixed plant material from three different organisms of the same group. For each sample, about 1 g of the frozen tissue was lyophilized in 50 ml falcon, and freeze-dried material was stored at −20C. Еach sample was prepared and analyzed in three replicates.

For the analysis, about 50 mg of lyophilized powder was weighed and treated with 7 ml 70% methanol for 30 min in an ultrasonic bath, then centrifuged for 10 min at 4,000 rpm. The supernatant was collected, filtered with Phenex GF/PVDF syringe filter (Ø 30 mm, 0.45 μm) and analysed on LCMS instrument. Analyses were performed by Shimadzu 8030 system consisting ofHPLC coupled to PDA and triple quadrupole mass spectrometer (HPLC-DAD-ESI-TQ MS). The chromatographic separation was performed on Discovery C18 column 4.6×150 mm, 5 μm in a gradient mode with mobile phase components A (0.3% acetic acid in water) and B (acetonitrile). The gradient run was performed in the following way: 0 – 4 min 10–40% B, 4 – 5 min 40-80%, 5 – 10.5 min, isocratic elution with 100% B, and then returned to the initial condition. The column temperature was 40C, the flow rate was 1 ml / min, the sample injection volume was 20 μL.

The ESI source was set in negative ionization mode. Multiple reaction monitoring was used to perform mass spectrometric quantification. MS conditions: interface voltage 3500V (ESI−), nebulizer gas (nitrogen) flow 2.5 l/min, drying gas (nitrogen) flow 15 l/min, CID gas pressure 60 kPa, DL temperature 250C, heat block temperature 400C. High purity argon was used as collision gas. The precursor and product ions (m/z) of target analytes were 178.95 and 134.95 for caffeic acid, 245.05 and 159.00 for hispidin; collision energy was 35V for both compounds.

Due to the lack of isotope-labeled standards, we added standards to samples to account for substantial matrix effect. Each sample was analysed twice, with and without the addition of standards. After the first analysis, a solution with known amount of caffeic acid and hispidin was added. Assuming a linear relation between the observed signal and concentration of compounds, concentration of the extract was calculated as C_extr_ = C_ad_ * S_extr_ / (S_tot_ − S_extr_), where C_ad_ – concentration of the added compound in the extract, S_extr_ and S_tot t_ – analyte peak area in the first and second analyses.

### Expression in cultured mammalian cells and luminescence imaging

DNA coding for RcTAL, HpaB, HpaC, nnHispS, NpgA, nnH3H, nnCPH, nnLuz was ordered synthetically (Evrogen, Russia) and cloned into the pKatushka2S-C1 vector (Evrogen) instead of Katushka2S coding sequence. HEK293T cell line was transfected with a mixture of all eight plasmids by FuGENE HD transfection reagent (Promega). Transfected cells were grown in DMEM medium (Paneco) supplemented with 10% fetal bovine serum (HyClone), 4 mM L-glutamine, 10 u ml−1 penicillin, 10 ug ml−1 streptomycin, at 37°C, 5% CO2. 24 hours after transfection, the medium was changed to MEM medium supplemented with 20mM HEPES, and luminescence was analysed by IVIS Spectrum CT (PerkinElmer). For the analysis, the background luminescence signal from the empty wells was subtracted from the luminescence signal of wells with control and autoluminescent cells.

## Data availability

The datasets generated or analysed during the current study are available from the corresponding authors on reasonable request.

## Acknowledgements

This study was designed, performed and funded by Planta LLC. We would like to thank Dr. Keith Wood for assisting in manuscript development. Planta acknowledges support from the Skolkovo Innovation Centre. We thank Milaboratory (milaboratory.com) for the access to computing and storage infrastructure. We thank Sergey Shakhov for providing photography equipment. The Synthetic biology Group is funded by the MRC London Institute of Medical Sciences (UKRI MC-A658-5QEA0). KSS is supported by Imperial College Research Fellowship. Experiments were partially carried out using the equipment provided by the Institute of Bioorganic Chemistry of the Russian Academy of Sciences Сore Facility (CKP IBCH; supported by Russian Ministry of Education and Science Grant RFMEFI62117X0018). FAK lab is supported by the ERC grant agreement 771209 — CharFL. KSS acknowledges the support by the president’s grant 075-15-2019-411. Assembly of some of the plasmids used in the study was supported by the Russian Science Foundation grant 19-74-10102. Chemical synthesis of luciferin and its precursors was partially supported by the Russian Science Foundation grant № 17-14-01169. LC-MS/MS analysis of extracts was supported by the Russian Science Foundation grant № 16-14-00052.

## Authors contributions

TM, ASM, LGS, TVC, EBG, TAK, NMM, SVC, AST, LIF, KAP, ESS, YVG, VVN, SAD, PVS, OAM, VOS, SMD, AIB, ASP, and KSS performed experiments. TM, ASM, LGS, TVC, EBG, TAK, NMM, SVC, AST, LIF, KAP, ESS, YVG, VVN, SAD, PVS, OAM, VOS, SMD, AIB, ASP, VVC, SVD, FAK, IVY and KSS performed data analysis. ASM designed imaging setup, planned and performed experiments, analysed data and wrote the paper. IVY and KSS proposed and directed the study, planned experimentation and wrote the paper. All authors reviewed and commented on the paper draft.

