## Supplementary text and figures for "Plants with self-sustained luminescence"

**Supplementary Table 1.** Concentration of caffeic acid and hispidin in leaves and flowers of *Nicotiana tabacum* plants measured by LC-MS/MS (μg/g of dry weight).

| Nicotiana tabacum line | Genotype | Caffeic acid in leaves at 3 am, μg/g | Caffeic acid in leaves at 3 pm, μg/g | Caffeic acid in flowers, μg/g | Hispidin in flowers *, μg/g |
|---|---|---|---|---|---|
| NT000 | *Wild type Nicotiana tabacum* | 2,51±0,46 | 2,32±0,25 | 0,76±0,004 | <0,05 |
| NT001 | nnHispS, nnH3H, nnLuz, nnCPH, KanR | 2,24±0,63 | 1,56±0,29 | 0,73±0,24 | 0,08±0,02 |
| NT078 | nnH3H-nnLuz fusion protein, KanR | no data | 1,80±0,12 | no data | no data |

* In leaves of *N. tabacum*, we also found trace amounts of hispidin, both in the wild type and transgenic lines. Hispidin was only detectable if we concentrated sample 20-fold with solid-phase extraction cartridges, however, this procedure negatively affected reproducibility of measurements.

** The content of caffeic acid was different in the wild type tobacco NT000 and transgenic line NT001 at 3 pm ($p \leq 0.05$, Student's t-test).

**Supplementary Table 2.** Primers used for plant genotyping

| Gene | Primer sequences | Amplicon size |
|---|---|---|
| nptII (KanR) | 5’- GCTATGACTGGGCACAACAGACAATC -3’<br>5’-TCCGAGTACGTGCTCGCTCGA -3’ | 381bp |
| nnLuz | 5’-CAATAGCATTCCCAATTATCCGAAGAG-3’<br>5’- ACAATCTTACCAGCAGGATCGTTAGTCA-3’ | 691bp |
| nnCPH | 5’-GTAGAGAAGGTAAGACAATTCAAGCATACGA -3’<br>5’-TCTTCTCGAACTGTATTTGCGAGAGTTC-3’ | 775bp |
| nnHispS | 5’-TGGATGTATTTCTCGACACGGCTAGA-3’<br>5’-TCAGCTCTGTCGGATATGTTGAAGGA -3’ | 735 bp |
| nnH3H | 5’-AGCATCAAAGGATGACTTGTTTCGAGT-3’<br>5’-GCTGAGTTAGAGCTCCTAAGCAAGGT-3’ | 465 bp |

#### Supplementary Note 1. Engineering of caffeic acid cycle pathway in plants.

*Neonothopanus nambi* caffeic acid cycle produces and metabolises 3-hydroxyhispidin from caffeic acid. The reactions are catalysed by four enzymes: luciferase nnLuz, hispidin synthase nnHispS, hispidin-3-hydroxylase nnH3H and oxyluciferin recycling enzyme nnCPH.

Since the 4'-phosphopantetheinyl transferase activity required for posttranslational modification of nnHispS [1] is likely present in tobacco [2,3], and caffeic acid is abundant in plants [4], we hypothesised that constitutive expression of only three enzymes may be sufficient for self-sustained bioluminescence and that further addition of the luciferin-recycling enzyme may increase the metabolic efficiency of the pathway.

Since residual amounts of hispidin were found in wild-type *N. tabacum* plant, we first created a plant line constitutively expressing fusion protein nnH3H-nnLuz. This strain did not emit light in a self-sustained manner but was luminescent upon injection of hispidin and luciferin.

Expression of three genes, *hisps*, *h3h* and *luz*, was sufficient to make tobacco plants autonomously bioluminescent. Additional expression of putative luciferin recycling enzyme nnCPH did not increase the brightness of plants suggesting the existence of a kinetic bottleneck at another enzymatic step in the cycle or a different function for the enzyme. For most experiments in this study we used the line NT001 containing two copies of the construct (**Supplementary Figure 10**).

#### Supplementary Note 2. Toxicity of caffeic acid cycle

The overall phenotype, leaf color, leaf size, flowering time, and seed germination of transgenic plants did not differ from those of the wild-type tobacco suggesting that unlike bacterial bioluminescent system [5], high expression of caffeic acid cycle is not toxic and does not impose a significant burden on plants (**Supplementary Fig. S1, S2**). We noticed that exposure to intense sunlight resulted in more abundant areas of necrosis in older leaves of transgenic plants in comparison to the wild-type plants. We also noticed that transgenic glowing plants were slightly taller than the wild type plants.

#### Supplementary Note 3. Activity and regulation of phenylpropanoid metabolism in plants.

Availability of phenolic acids in plants is controlled through expression of genes of central phenylpropanoid pathway coding for phenylalanine ammonia-lyase, cinnamic acid 4-hydroxylase and p-coumaric acid 3-hydroxylase [6,7], as well as by expression of enzymes that use phenolic acids as a substrate (**Figure 1**). The phenylpropanoid metabolism is additionally shaped by tissue-specific distribution of numerous isoforms of these enzymes, compartmentalization, metabolic channeling and expression of membrane transporters [8]. Enzymes of phenylpropanoid pathway are typically associated in metabolons to increase reaction rates, localise toxic intermediates and channel metabolism towards different competing branches of biosynthesis [8]. In central phenylpropanoid metabolism, phenylalanine ammonia-lyase colocalizes with cinnamic acid 4-hydroxylase and 4-coumarate:CoA ligase on the cytosolic side of the endoplasmic reticulum membrane [8,9] limiting leakage of intermediates into the cytosol.

Steady-state concentration of luciferin in glowing plants should depend on metabolic flux through phenylpropanoid metabolism being limited by the concentration of caffeic acid available to HispS. Production of caffeic acid reflects the activity of the central phenylpropanoid metabolism [10]. Commitment to this pathway is controlled by phenylalanine ammonia-lyase. Its promoters exhibit developmental control and are active in roots, in particular, in vascular and endodermal tissues of lateral roots, leaf and petal tips, pigmented regions of petals and stamen filaments [11,12]. Similarly, expression profiles of C4H and C3H (CYP98A3) include anthers, apical part of pistils and lignifying tissues: leaf veins, stem vascular bundles and root central cylinder [13]. High level of isoforms of 4-Coumarate:CoA Ligase (4CL) in *Arabidopsis* was also revealed in lignified tissues. 4CL4 was reported to be expressed in the epidermis, cortex, cambium, phloem, and pith. 4CL1 and 4CL3 activity was detected in flowers and roots [14]. These patterns overlap with the observed distribution of luminescence in glowing plants (**Figure 2, Supplementary Figures 4, 9**)

**Supplementary Note 4. Can hispidin precursors pass through cell membranes?**

Hispidin biosynthesis is a multi-component multistep reaction that requires caffeic acid, malonyl-CoA and ATP.

Cell membranes are not passively permeable to coenzyme A derivatives. It is not clear whether plant cells can actively uptake malonyl-CoA from the environment.

Another component of the reaction is caffeic acid. Plant cells are able to uptake phenolic acids [15] and amino acids [16] from the environment.

Another compound, ATP, cannot passively cross the membranes. Tracer experiments confirmed influx of externally applied ATP into plant cells [17] but extracellular ATP is also known to act as a signalling molecule in plants [18].

In addition, there is a negative feedback in phenolic metabolism: caffeic acid is known as PAL inhibitor [19]. After infiltration with caffeic acid, its biosynthesis may be arrested until the excess of caffeic acid is metabolised, explaining why intensity of luminescence at the sites of injection drops slightly below the initial level before injection.

#### Supplementary Note 5 on circadian oscillations

*nnluz* gene was expressed at high levels throughout the day, with time-dependent oscillations (**Supplementary Figure 11**). The expression decreased during the night (21:00 to 6:00) and increased after the sunrise (05:45 on the day of collection).

Experiments with continuous night (**Figure 3c, 3d**) revealed two peaks of luminescence in complete darkness, corresponding to the subjective day of plants. This indicates that the luminescence is controlled by the circadian clock in tobacco. The subjective time also affects nyctinastic leaf movements as estimated by oscillations of the top line on **Figure 3c**.

The subsequent two dark 24-hour periods lack any circadian oscillation of luminescence, correlated to nyctinastic movement deprivation. These observations indicate that in tobacco plants the circadian clock is effectively switched off on the third and fourth 24-hour periods in the dark.

The circadian rhythm of both luminescence and nyctinastic leaf movements restores completely within two days of normal lightening (**Figure 3c, 3d**).

### Supplementary Figures

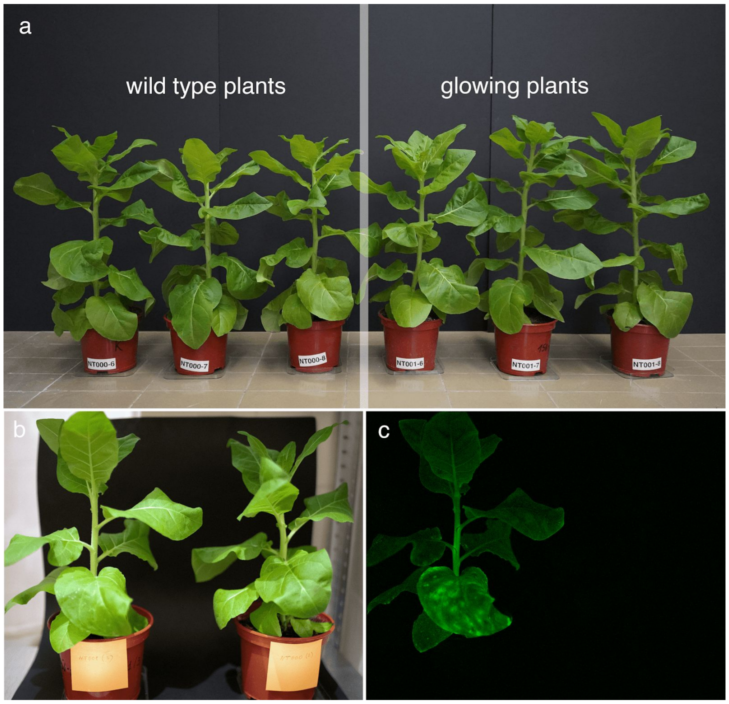


**Supplementary Figure 1.** Phenotype of glowing and wild type *Nicotiana tabacum* plants. Potted glowing plants and wild type plants of the same age in ambient light (A, B) and in the dark (C).

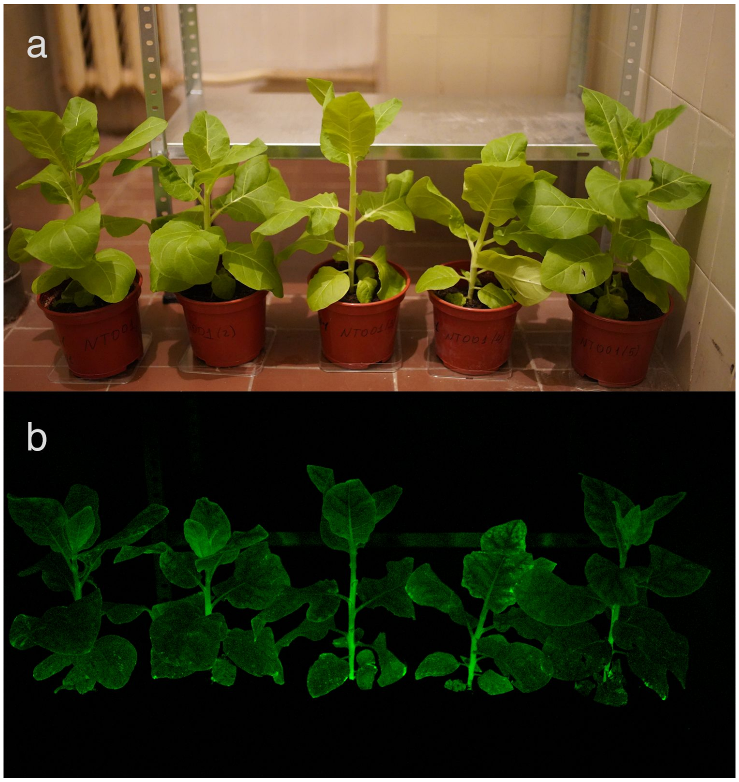


**Supplementary Figure 2.** Six-week glowing plants in ambient light (A) and in the dark (B).

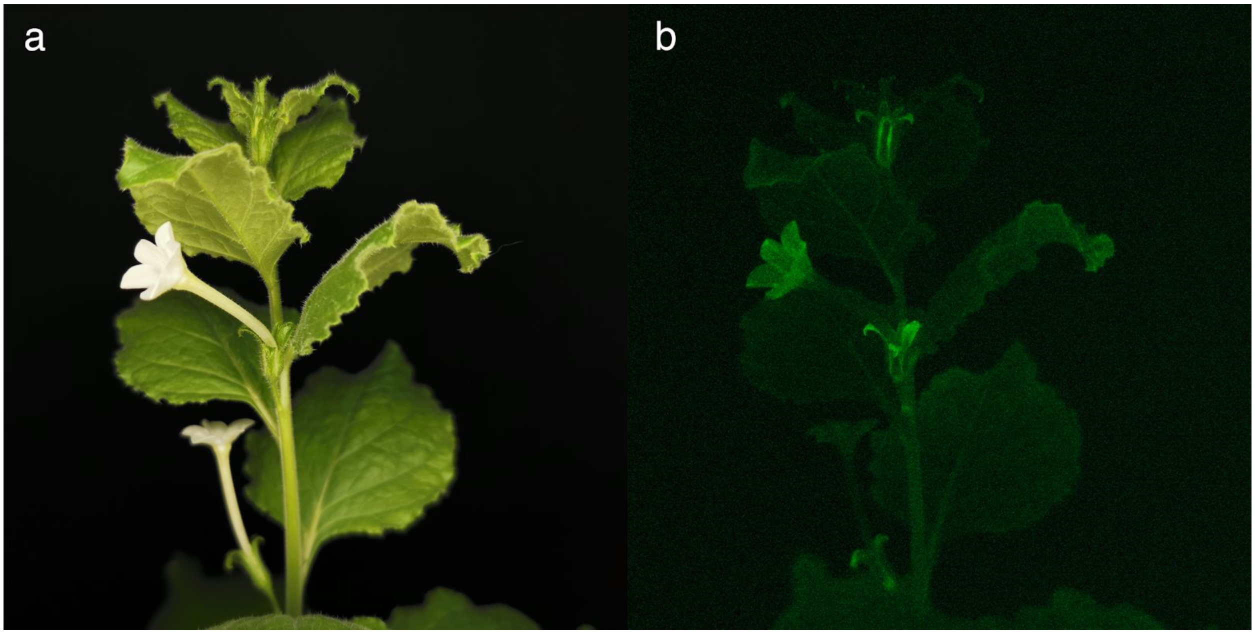


**Supplementary Figure 3.** Photos of glowing *Nicotiana benthamiana* plant taken on a smartphone in ambient light (A) and in the dark with 30-second exposure (B).

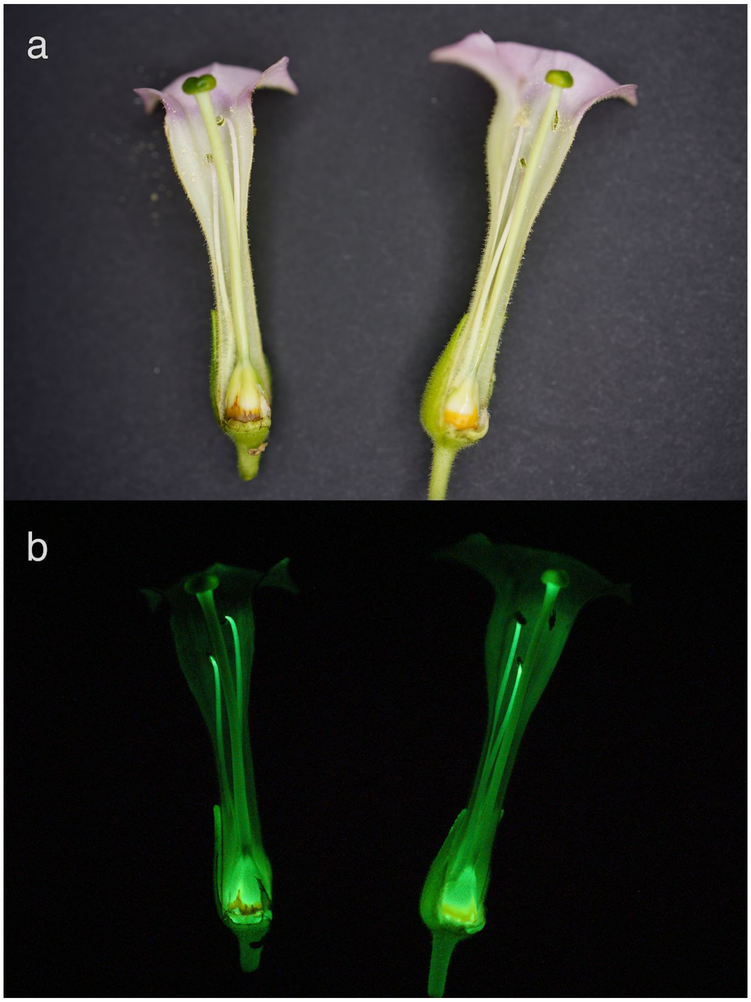


**Supplementary Figure 4.** Photos of cross-section of flowers from a glowing *Nicotiana tabacum* plant in ambient light (A) and in the dark (B).

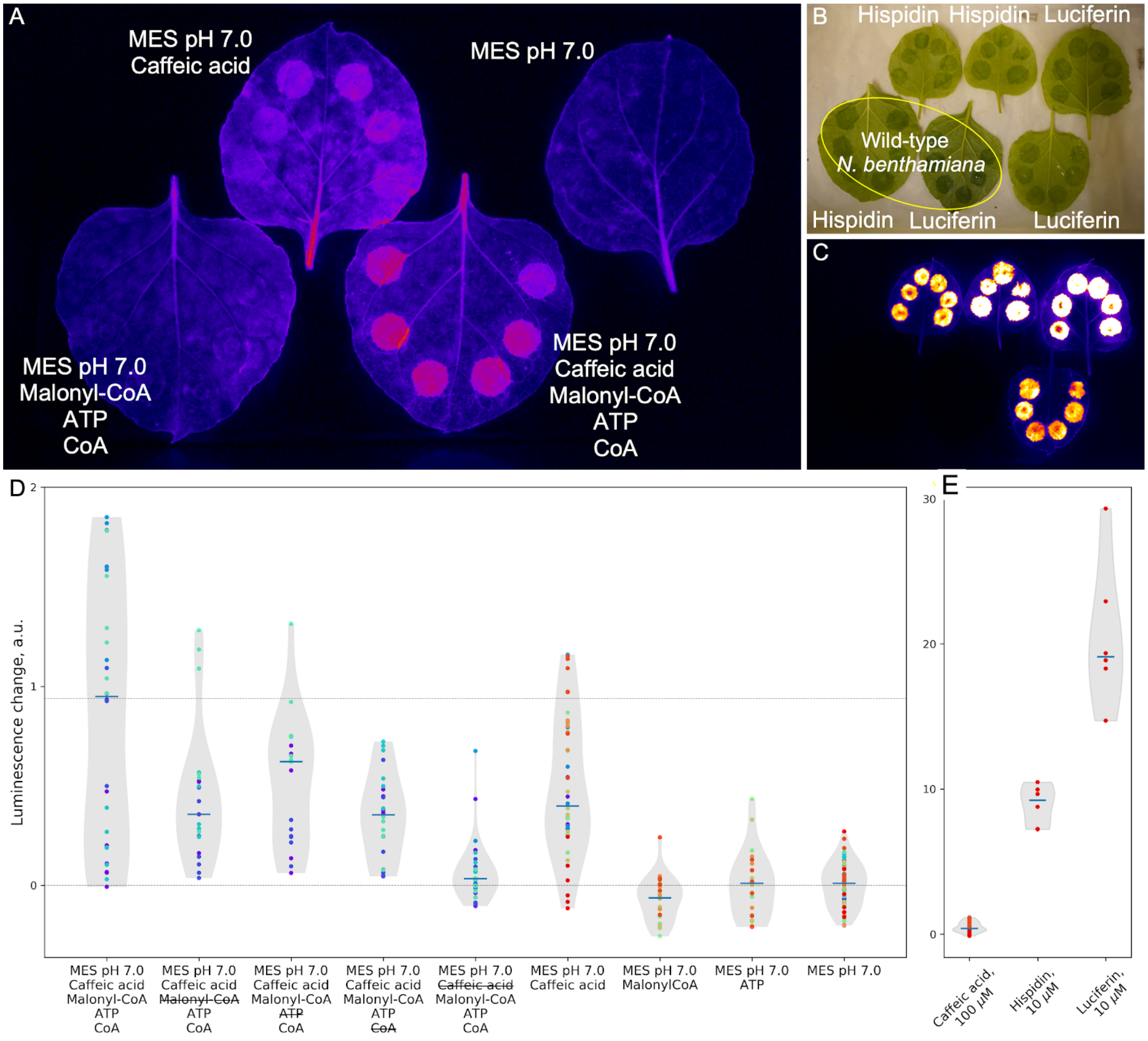


**Supplementary Figure 5.** Infiltration of *Nicotiana benthamiana* leaves with solution of hispidin precursors. **A.** Representative photo of transgenic bioluminescent *Nicotiana benthamiana* leaves injected with solution of individual hispidin precursors. **B, C.** Injections of hispidin and luciferin in leaves of bioluminescent and wild-type *N. benthamiana* plants. **D.** Violin plots showing luminescence response at sites of injections of mixtures of hispidin precursors of various composition. Data points are shown, colored according to the leaf of origin. Blue horizontal line represents median. Luminescence change is absolute change of signal (ΔL = luminescence after injection – initial luminescence). Strikethrough text highlights absence of a compound in the mixture. Notably, only mixtures containing caffeic acid induced the increase of luminescence. **E.** Effect of injections of three successive luciferin precursors on the intensity of luminescence of *N. benthamiana* leaves.

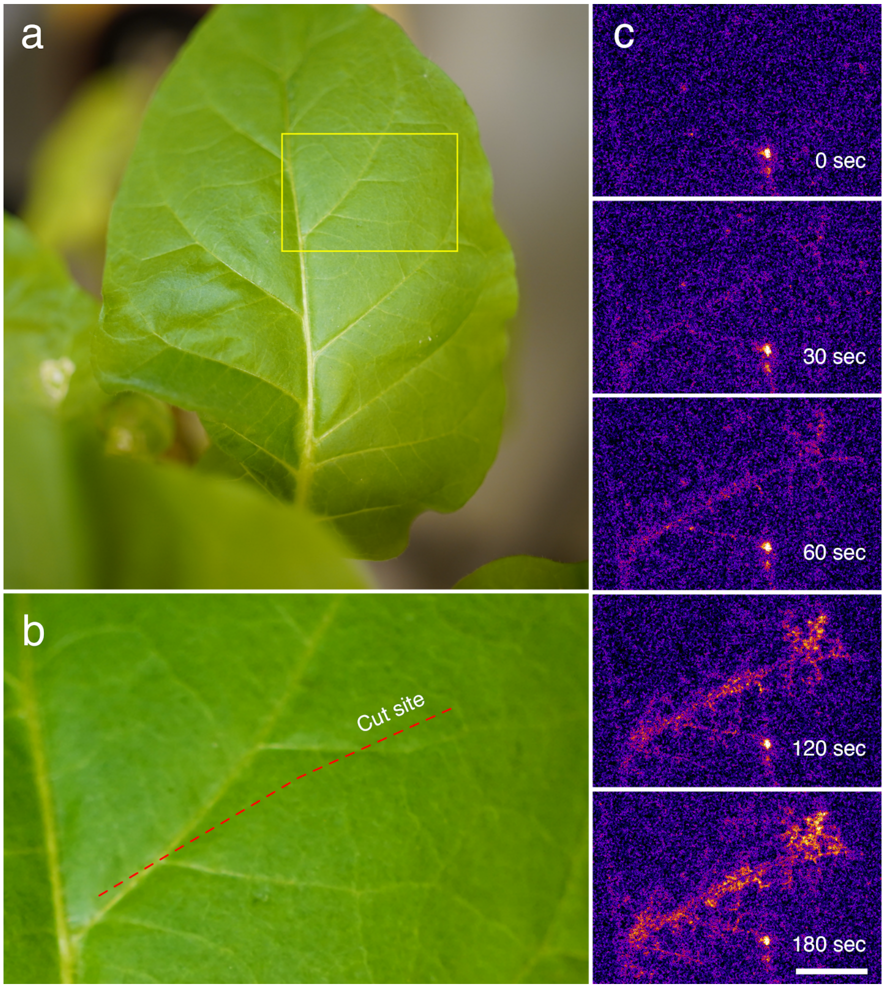


**Supplementary Figure 6.** Dynamics of bioluminescence following injuries in leaves.
**A.** Representative leaf before injury. **B.** Close-up photo of the cut site. **C.** Dynamics of bioluminescence at the cut site following injury (exposure time — 5 seconds). Scale bar corresponds to 1 cm.

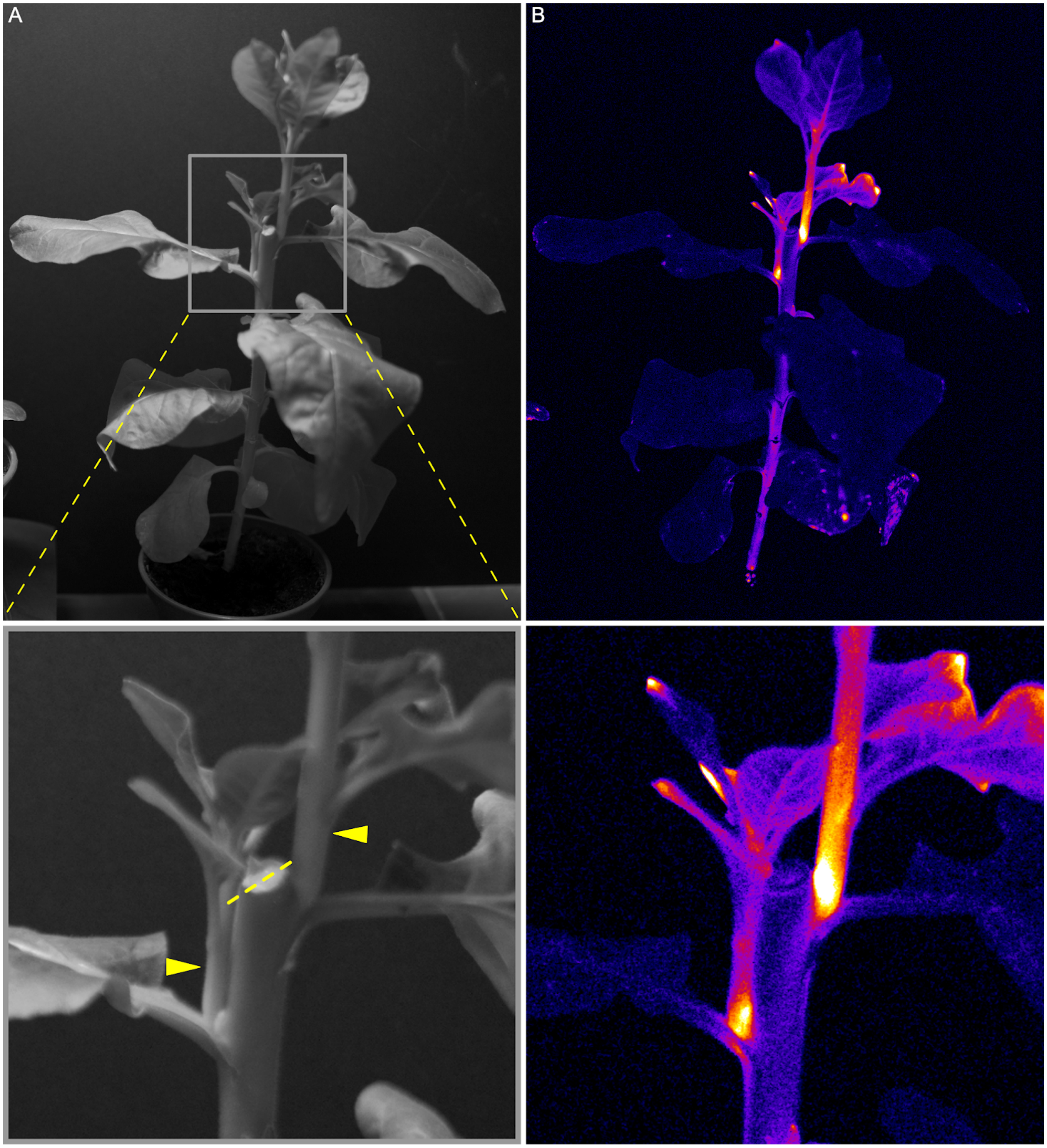


**Supplementary Figure 7.** Pruning-induced sustained increase in bioluminescence in lateral shoots.
**A.** Photo under ambient light. Lateral shoots are marked with yellow arrows. The cut is highlighted by dashed yellow line on the enlarged photo (bottom). **B.** Photo captured in the dark.

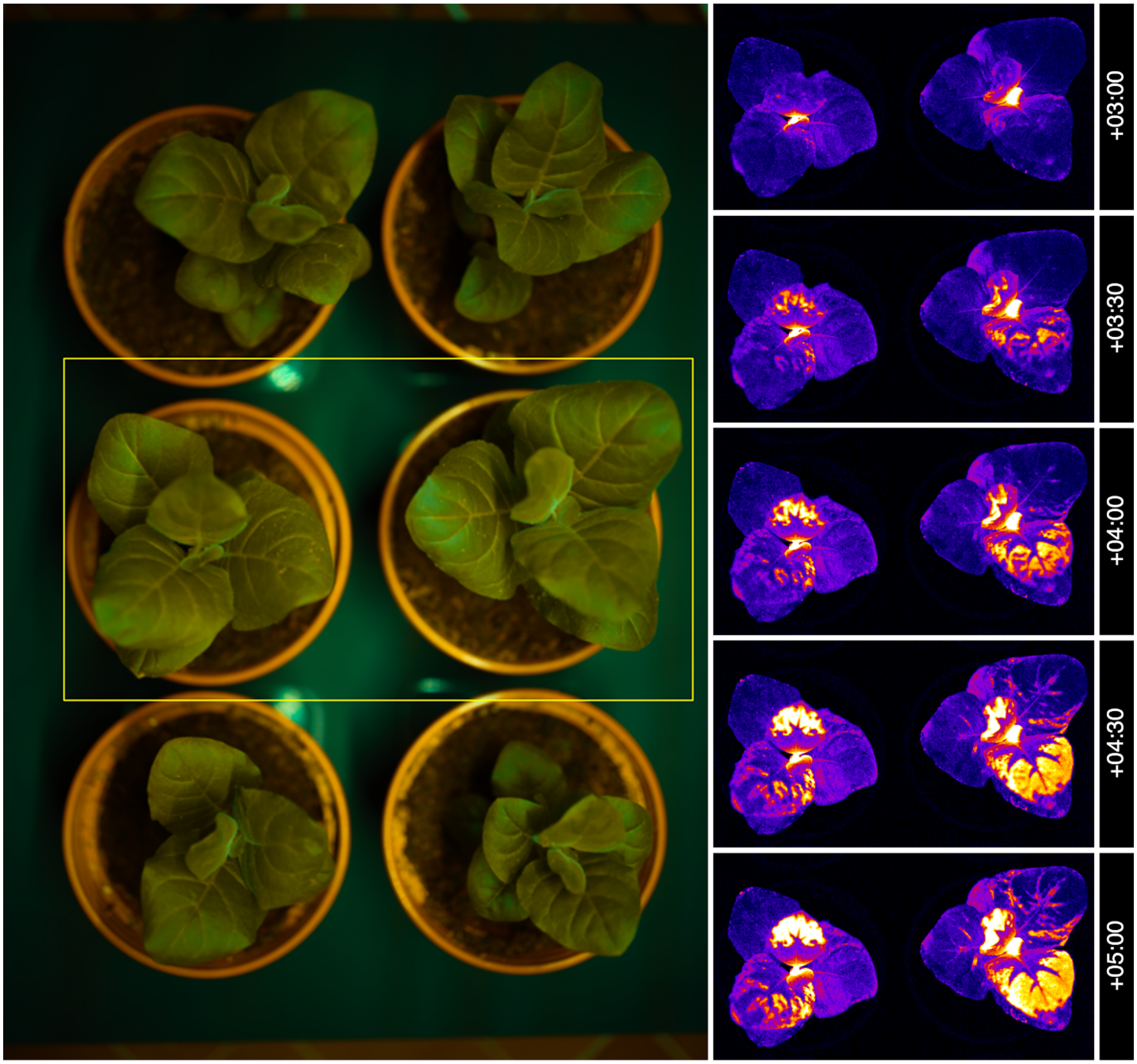


**Supplementary Figure 8.** Dynamics patterns of bioluminescence in leaves of young glowing plants. Time relative to the start of the day is designated on the right (HH:MM format).

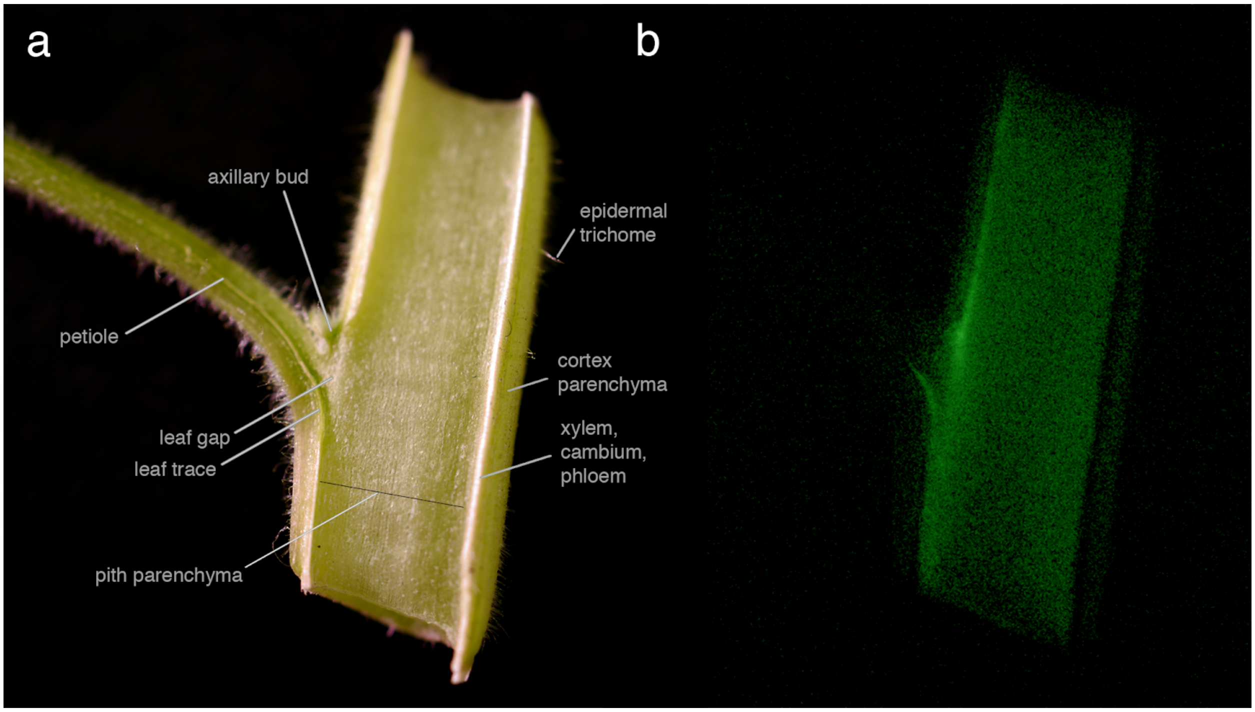


**Supplementary Figure 9.** Photos of cross-section of stem of a glowing *Nicotiana tabacum* plant in ambient light (A) and in the dark (B). The robust light emission belongs to the cells of shoot pith parenchyma (in the center of the stem). Weaker bioluminescence is observed from core parenchyma cells (periphery of the stem). Maximal light intensity corresponds to the region of the axillary bud — newly forming meristem — and surrounding parenchymatous cells. Completely lignified tissues, xylem and phloem, with cambium in between, lack luminescence (visible at the right side of the stem as a dark band).

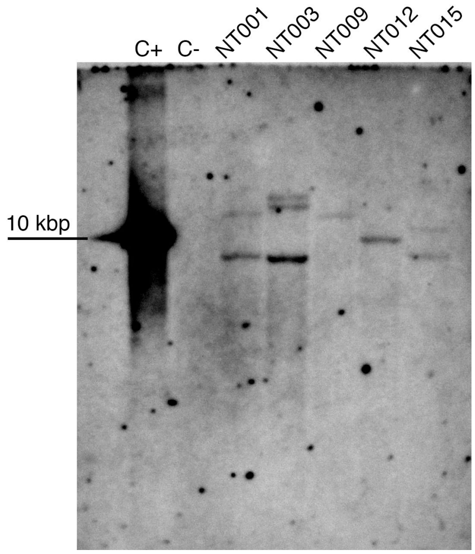


**Supplementary Figure 10.** Southern blot of DNA extracted from various glowing transgenic tobacco lines, with the probe annealing to *nnluz* gene. There are two copies of nnLuz gene in the genome of the line NT001 (used for all *N. tabacum* experiments in this study).

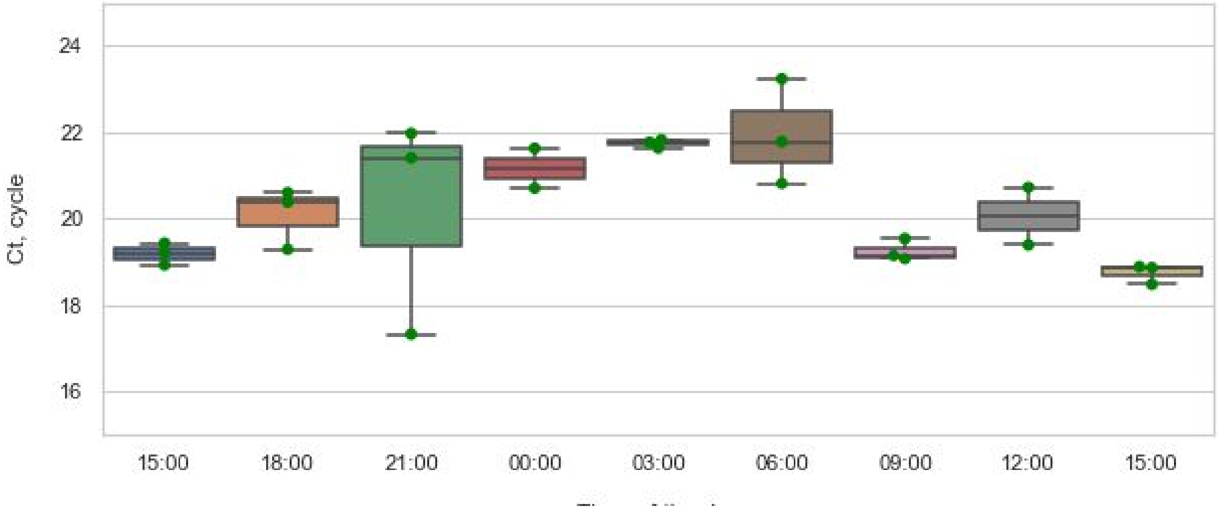


**Supplementary Figure 11.** Expression of nnLuz in *N. tabacum* throughout the day measured by qPCR. Plants were grown under natural light conditions. Each data point represents a different biological sample (measured in three technical replicates, the mean value of technical replicates is shown). Error bars represent standard deviation.

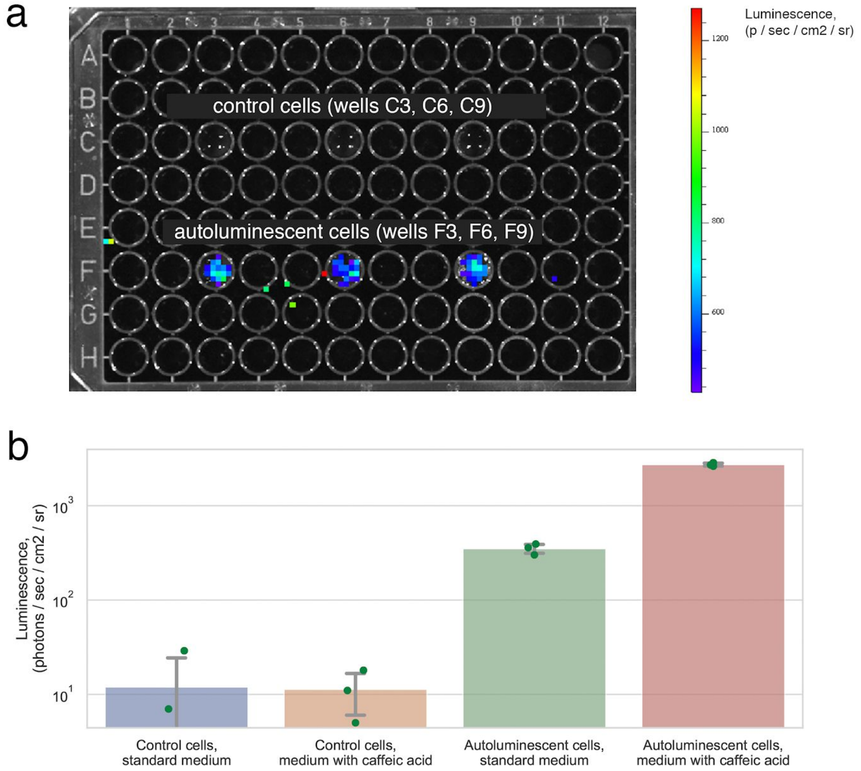


**Supplementary Figure 12. Engineering of autonomous luminescence in mammalian cells.**

**A.** Overlay of ambient light (black and white) and luminescence (pseudocolor) images of a multiwell plate with HEK293T cells co-transfected with plasmids encoding genes of fungal bioluminescence pathway and caffeic acid biosynthesis: RcTAL, HpaB, HpaC, nnHispS, NpgA, nnH3H, nnCPH, nnLuz.

**B.** Light emission from autoluminescent cells, in standard MEM medium with 20mM HEPES and in the same medium supplemented with caffeic acid (N=3, error bars represent standard deviation).

#### Supplementary files legends

**Supplementary Video 1. Germinating T1 seeds**

Time-lapse imaging of transgenic *N. tabacum* T1 seeds germinating in transparent agar. Overlay of photos in ambient light (grey) and luminescence (pseudocolor) is shown.

**Supplementary Video 2. Regenerating roots**

Time-lapse luminescence imaging of regenerating roots of cut shoots of transgenic *N. tabacum* plants (photos every 5 min). Luminescence intensity is shown in pseudocolor.

**Supplementary Video 3. Root microscopy**

Time-lapse microscopy luminescence imaging of regenerating roots of cut shoots of transgenic *N. tabacum* plants (photos every 5 min). Luminescence intensity is shown in pseudocolor.

**Supplementary Video 4. Long time-lapse throughout lifespan of plants**

Time-lapse imaging of transgenic *N. tabacum* plants from vegetation to flowering. Luminescence intensity is shown in pseudocolor. Yellow asterisks indicate daytime.

**Supplementary Video 5. Injection of hispidin in leaves of autonomously luminescent N. tabacum**

Time-lapse luminescence imaging following injection of hispidin (into the central area of the upper part of the blade) and caffeic acid (into the apical part) of leaves of autonomously luminescent *N. tabacum*. Luminescence intensity is shown in pseudocolor.

**Supplementary Video 6. Leaf injuries in N. tabacum**

Time-lapse luminescence imaging following injury of leaf blade of autonomously luminescent *N. tabacum*. Luminescence intensity is shown in pseudocolor.

**Supplementary Video 7. Dynamic luminescence in leaves**

Time-lapse luminescence imaging of pruning-induced lateral shoots of autonomously luminescent *N. tabacum.* Luminescence intensity is shown in pseudocolor.

**Supplementary Video 8. Time-lapse imaging of flowers**

Time-lapse luminescence of flowers of autonomously luminescent *N. tabacum*. Luminescence intensity is shown in pseudocolor.

**Supplementary Video 9. Dynamic patterns of luminescence ini leaves of young plants**

Dynamics patterns of bioluminescence in leaves of young glowing plants. Time relative to the start of the day (+HH format) as well as absolute time (HH:MM format) is displayed in the lower left corner of the video.
